# Functional characterization of 3D-protein structures informed by human genetic diversity

**DOI:** 10.1101/182287

**Authors:** Michael Hicks, Istvan Bartha, Julia di Iulio, Ruben Abagyan, J. Craig Venter, Amalio Telenti

## Abstract

Sequence variation data of the human proteome can be used to analyze 3-dimensional (3D) protein structures to derive functional insights. We used genetic variant data from nearly 150,000 individuals to analyze 3D positional conservation in 4,390 protein structures using 481,708 missense and 264,257 synonymous variants. Sixty percent of protein structures harbor at least one intolerant 3D site as defined by significant depletion of observed over expected missense variation. We established an Angstrom-scale distribution of annotated pathogenic missense variants and showed that they accumulate in proximity to the most intolerant 3D sites. Structural intolerance data correlated with experimental functional read-outs *in vitro*. The 3D structural intolerance analysis revealed characteristic features of ligand binding pockets, orthosteric and allosteric sites. The identification of novel functional 3D sites based on human genetic data helps to validate, rank or predict drug target binding sites *in vivo*.

---

Recent large-scale sequencing projects of the human genome and exome detail the extent of genetic diversity in the human population ^1-3^. To date, there are over 4.5 million amino acid-changing (missense) variants reported in the human exome. Much attention has been directed to the association of variants with disease ^3,4^. However, these data also represent an unprecedented opportunity to characterize protein structure-function relationships *in vivo*. In particular, the pattern of distribution of genetic variants describes the functional limits to structural and functional modifications for a given protein. Inference of critical three dimensional (3D) sites would also be informative for drug development and mechanisms of action, including selectivity, lack of response, or toxicity.

Finding important 3D sites within these structures has been done through a variety of methods. Genetics-based scoring metrics can measure the deleteriousness of genetic variants in a protein, a property that strongly correlates with both molecular functionality and pathogenicity ^5,6^. Scores may also consider interspecies conservation ^7^ to discover “constrained elements” indicative of putative functional elements. Recent sequencing efforts of human genomes and exomes provide a different level of spatial information through the saturation of proteins structures to derive human-specific intolerant sites ^3,6,8^. Previous approaches have emphasized gene level features (eg, essentiality, burden of variation), and linear analyses of variation in a gene rather than the distribution of variants in 3D space. However, additional methods that describe mutational clustering in protein structures have been created. Ryslik et al. described *iPAC* (identification of Protein Amino acid Clustering)^9^, SpacePAC (Spatial Protein Amino acid Clustering)^10^, *GraphPAC* (Graph Protein Amino acid Clustering)^11^, and *QuartPAC* (Quaternary Protein Amino acid Clustering)^12^. Fujimoto et al.^13^, Tokheim et al.^14^ and Meyer et al.^15^ analyzed 3D position and clustering of mutations using exome sequence data from The Cancer Genome Atlas (TCGA) from up to 7,215 samples and 23 types of cancer and over 975,000 somatic mutations. A comparison of algorithms for the detection of cancer drivers at subgene resolution has just been published ^16^.

It should be noted that scoring methods in oncology emphasize mutational clustering – as critically relevant in cancer biology – and not intolerance to variation in the human proteome at large. Most studies that analyze the relationship between point mutations and experimentally observed 3D protein structures published to date have been limited to individual proteins. Bhattacharya et al. ^17^ manually analyzed one single nucleotide variant in each of 374 human protein structures to assess the effects of genetic variation on structure, function, stability and binding properties of the proteins. Arodz et al. analyzed a limited set of pairs of proteins of the same length differing by a single amino acid ^18^. A thorough analysis of the proteome requires a larger study population to have sufficient saturation of the 3D space (ie. use of population genetics to determine the spatial distribution of variation, and the patterns of constraint as reflected in intolerance and tolerance to variation). Thus, we initiated a study that uses human genetic variation from nearly 150,000 human exomes and genomes and 4,390 x-ray protein structures to model tolerance to amino acid changes in the 3D space of the human proteome.

To understand variation in the structural proteome, we first identified 26,593 structures associated with 4,390 Uniprot entries that fulfilled our inclusion criteria: x-ray crystal structures with a defined resolution, a minimum chain length greater than 10 amino acids and at least 80% identity between the aligned matches of the Uniprot canonical sequence and the PDB structure. Given the multiplicity of possible structures for the 4,390 proteins, we chose as representative the structure with the most scored Uniprot features. In total, we mapped 139,535 Uniprot features to the structures, and extracted a 3-dimensional context by defining a 5-Angstrom radius space for each feature; hereafter referred to as a “3D-site”. We identified 481,708 missense variants for these proteins from the analysis of 146,426 individuals’ exomes. From these contextualized data, we constructed a model that describes functional constraints in three-dimensional protein structures (Figure 1, Methods section). The strength of constraint (intolerance) was reflected in a 3-dimensional tolerance score (3DTS) that summarized the differences between observation and expectation in genetic variation at the level of 3D-sites.

**Figure 1.**
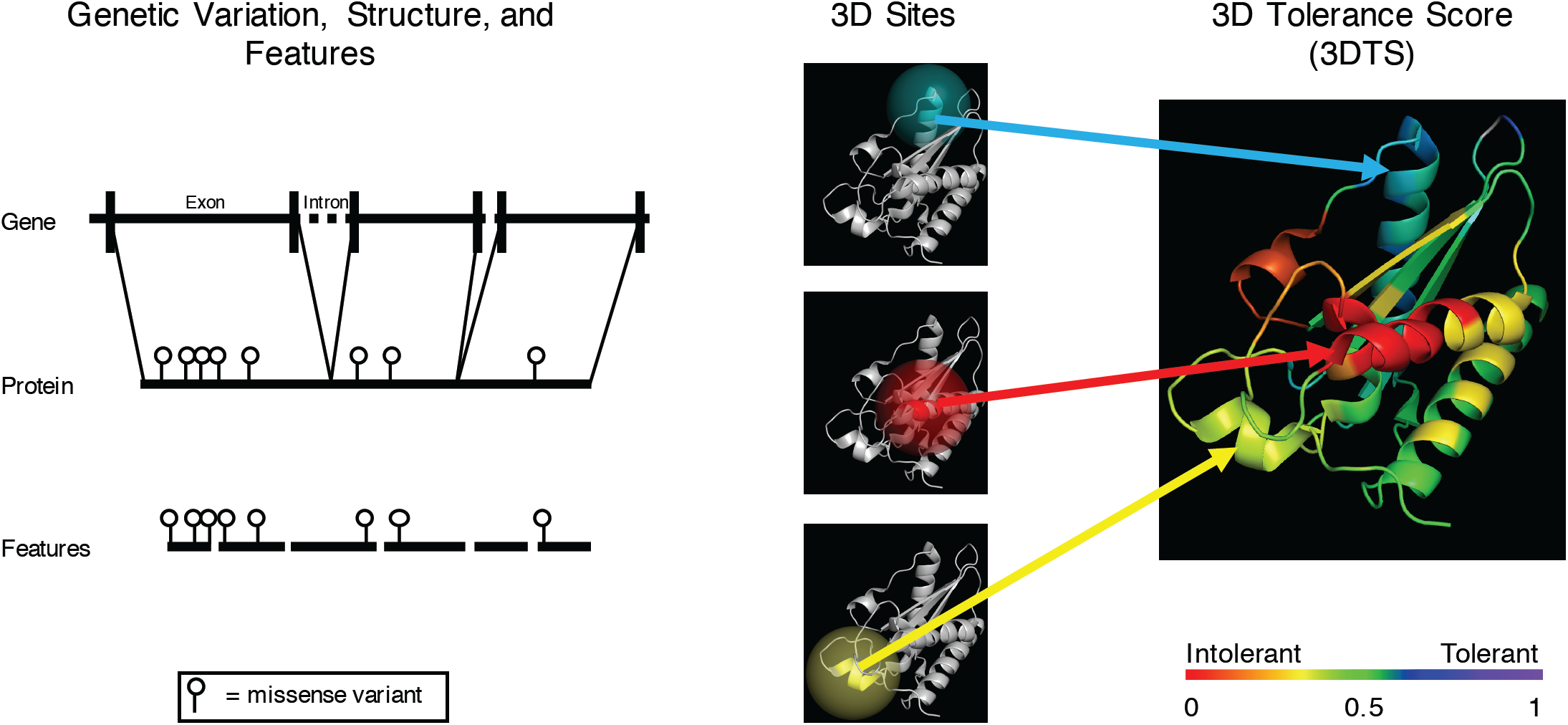
3D tolerance to variation in the proteome. Missense variation data from genome and exome sequencing projects are mapped to 3D protein structures. Features extracted from Uniprot are also mapped to the 3D structures. Using these features as reference points, a 3D context is constructed and the corresponding genetic data are extracted. A 3D tolerance score (3DTS) is generated from this information. The 3DTS values can be ranked and the corresponding tolerance ranks (or scores) can be projected back onto the 3D structure.

For the representative set of structures for the 4,390 proteins, we described the distribution of 3DTS values in Figure 2A. In total, 2,642 proteins had at least one intolerant 3D-site defined at the 20^th^ percentile (3DTS=0.33, approximately 70% depletion of observed over expected missense variation). The most intolerant 3D-sites corresponded to DNA binding sites, zinc fingers, and cross-linkages, while the most tolerant 3D-sites included transit peptides, non-standard residues (i.e., selenocysteines), and propeptides. Structural features (helix, turn, strand) showed median 3DTS values close to the median proteome-wide (Figure 2B). The precise interpretation of 3DTS values required the assessment of functional consequences of amino acid changes in intolerant versus tolerant 3D-sites. However, a challenge of functional testing proteome-wide is the requirement of cellular assays that are disease and gene relevant, robust, and scalable – a serious limitation that explains that to this date, the experimental characterization of all possible missense variants in a mammalian gene has been limited to one full protein, PPARG ^19^, and two single protein domains of BRCA1 (the RING domain) and YAP65 (the WW domain) ^20,21^. We therefore sought to validate 3DTS against the available functional data for these proteins and domains. In the case of the WW domain in YAP65, positional functional data were not easily accessible and the domain represented a set of only 25 amino acid positions; therefore, it was not assessed.

**Figure 2.**
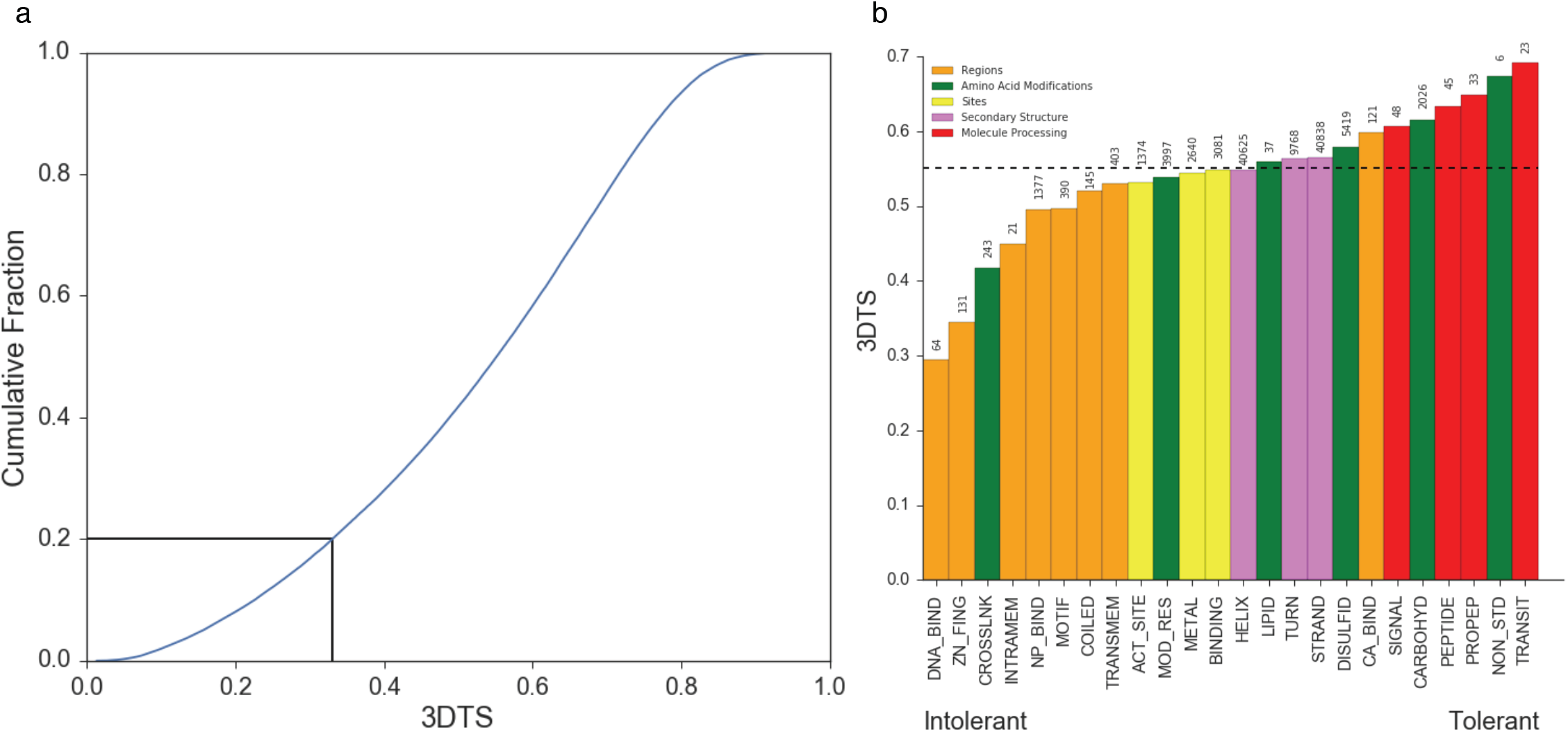
Distribution of tolerance values across the structural proteome. (a) Distribution of 3DTS values for 139,535 3D-sites for structures representing 4390 proteins. The 3DTS value at the 20^th^ percentile (3DTS < 0.33), is used to define intolerant sites. (b) Median 3DTS for a subset of feature types. The number of each feature type with a 3DTS value is shown above each column. The overall median across the structural proteome is represented by a horizontal dashed line. Feature types are colored by Subsections defined by Uniprot (http://www.uniprot.org/help/sequence_annotation).

PPARG is a drug target for thiazolidinediones and newer partial PPARG modulators used in the treatment of diabetes ^22^. PPARG exemplifies the challenge of classifying newly identified variants even in a well-studied protein implicated in disease. In the original work ^19^, functional interpretation of PPARG variants required the construction of a cDNA library consisting of all possible amino acid substitutions in the protein. The library was introduced into human macrophages edited to lack the endogenous PPARG, and stimulated with PPARG agonists to trigger the expression of CD36, a canonical target of PPARG. Sorted CD36+ and CD36- cell populations were sequenced to determine the distribution of each PPARG variant in relation to CD36 activity. We showed a strong correlation (r^2^=0.47, p=0.0001) between the 3D- sites defined by 3DTS and the functional scores described in Majithia et. al. ^19^ Specifically, both the *in vitro* and the *in silico* scores identified the DNA-binding and ligand binding sites as intolerant to missense variation, while the hinge domain reflected increased tolerance to missense variation (Figure 3). The 5 Å context also showed stronger correlations than the linear features (a 0 Å context), a 3 Å context or a 7 Å context (Supplementary Figure S1). Additionally, Majithia et al. indicated that their transgene library may not have detected all possible functional effects of coding variation, suggesting that the concordance of r^2^=0.47 between *in vitro* and *in silico* readouts should be interpreted as conservative ^19^.

**Figure 3.**
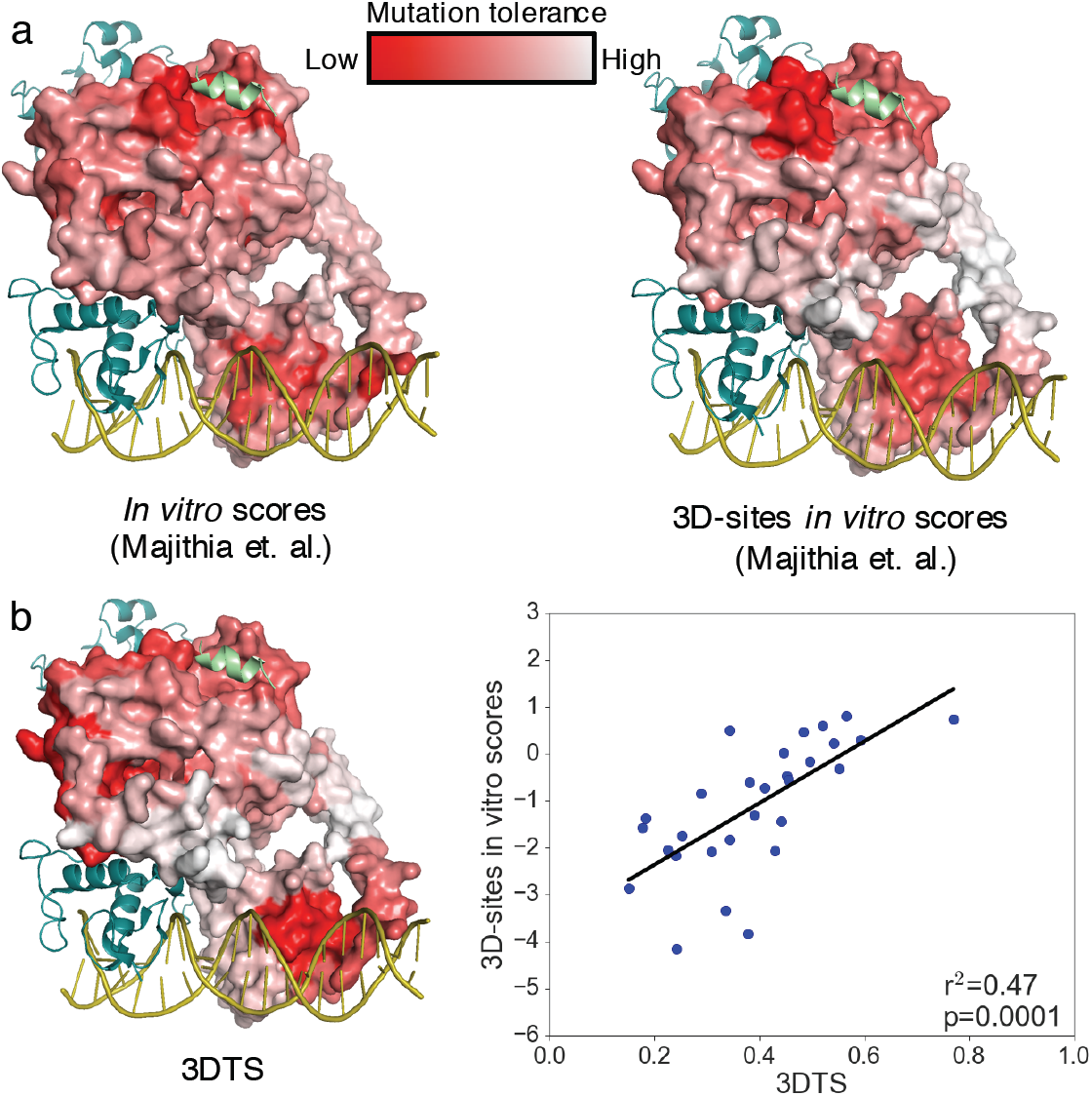
Comparison of *in vitro* functional data and *in silico* data for the DNA binding and ligand binding domains of PPARG. (a) Projection of the integrated functional scores described in Majithia et. al.^19^ for each amino acid, and the scores averaged across the 3DTS-defined sites for a crystal structure 3dzy ^32^. The color scheme is chosen to match the one described in Majithia et. al. (b) A projection of 3DTS onto PPARG is seen on the left and the 3D-site level correlation between 3DTS and the 3D-site averaged *in vitro* scores is shown in the plot on the right.

The second example uses BRCA1; an informative exercise because the approach is validated for only one of the structural domains (RING). The RING domain represents only 5% of the canonical BRCA1 protein; however, 58% of the pathogenic missense substitutions occur within this domain ^20^. In the original work, functional analysis of the RING domain required testing for two functions: BRCA1 E3 ligase activity in phage display assays, and interaction with BARD1 in yeast two-hybrid assays ^20^. The combination of these two molecular functions into a larger biological function, homology directed repair, resulted in a concordance of r^2^=0.32, p=0.033, with the rank values of 3DTS. The zinc-binding sites showed the greatest intolerance in this structure (Figure 4). In summary, the *in silico* 3DTS values recapitulate *in vitro* functional data without engaging in complex assays that require extensive and time-consuming *in vitro* assays and dedicated functional readouts.

**Figure 4.**
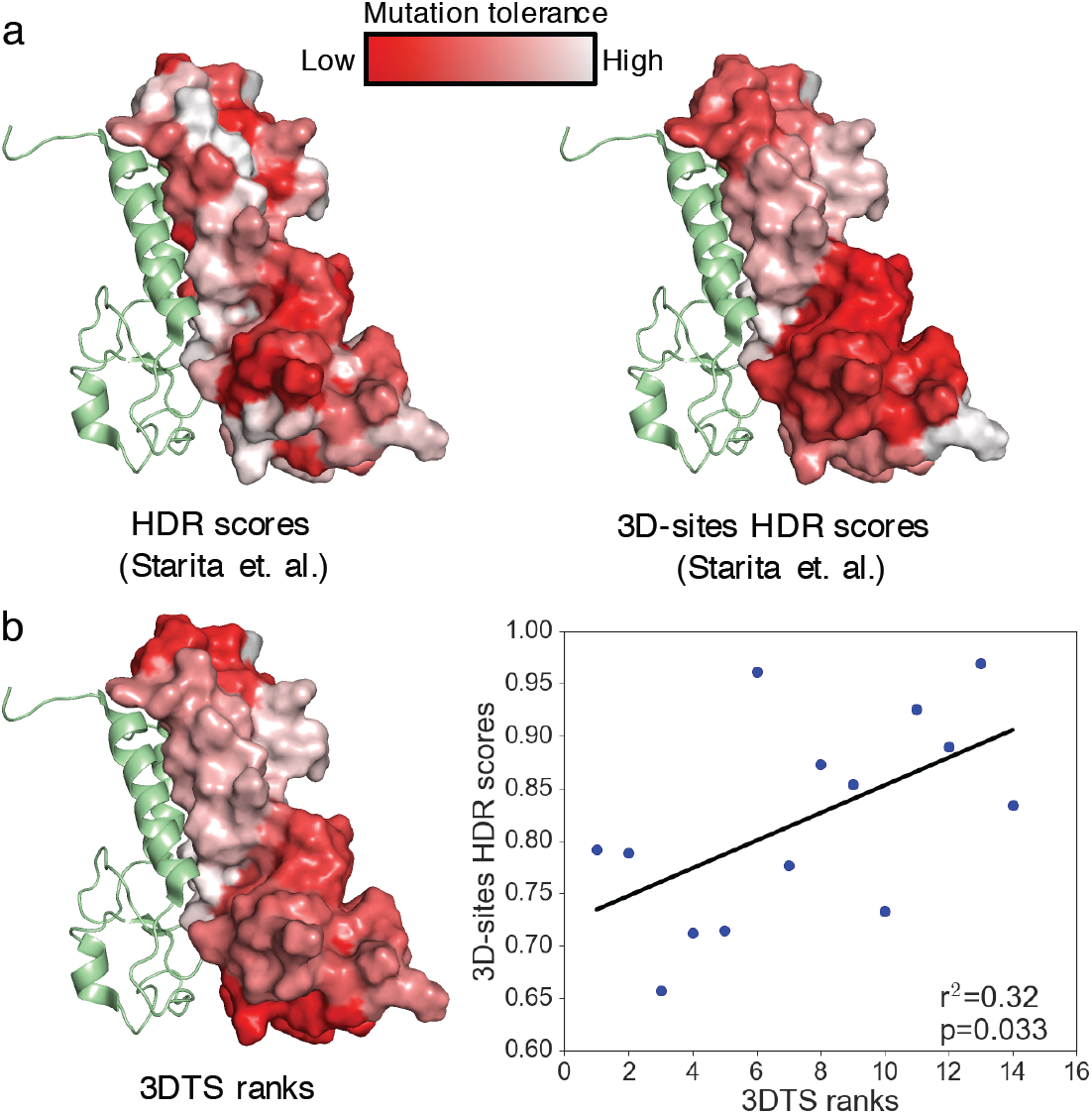
Comparison of *in vitro* functional data and *in silico* data for the RING domain of BRCA1. (a) Projection of the homology directed repair (HDR) scores described in Starita et al.^20^ averaged across amino acids, and averaged across the 3DTS-defined sites. (b) A projection of 3DTS ranks onto the RING domain of BRCA1 is seen on the left and the 3D-site level correlation between 3DTS rank and the 3D-site averaged HDR scores is seen in the plot on the right.

Predicting functionally intolerant 3D sites, and the distribution of variants with respect to these sites, may have several practical applications. For example, variants within intolerant sites may carry phenotypic consequences (i.e., pathogenicity). We thus aimed at establishing the association between 3D intolerance to variation and pathogenicity of variants. We identified 192 structures with at least one pathogenic missense variant (3081 total variants) and at least one common (allele frequency > 1%) missense variant (373 total variants). The distance between the closest atoms of the most intolerant feature and each variant were measured. In this set, the greatest enrichment of pathogenic relative to common variants appeared within the most intolerant site (2.3-fold enrichment) and another peak in enrichment was seen within ∼6-14 Å of the most intolerant site (Figure 5, Supplementary Figure S2). Due to the scarcity of common missense variants, we also used synonymous variants as a proxy for neutral variation, which increased the number of available structures to 438 and the number of pathogenic missense variants to 9,531, leveraging a total of 26,229 synonymous variants. In this set, the greatest enrichment of pathogenic variants was observed ∼4-9 Å away from the most intolerant site (Figure 5, Supplementary Figure S2). The enrichment of pathogenic variation diminishes with distance.

**Figure 5.**
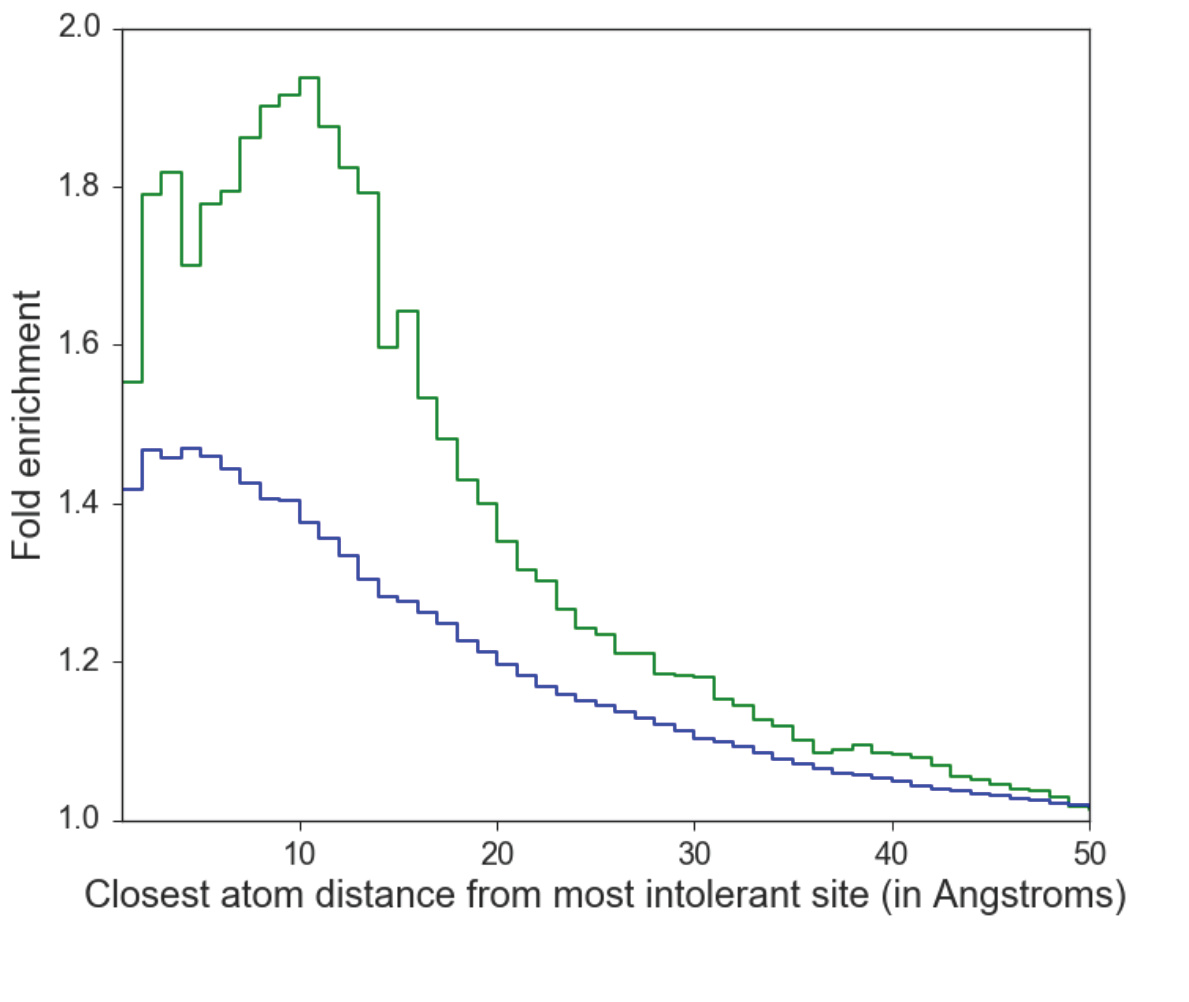
Relationship between tolerance and pathogenicity. Distance mapping of pathogenic variants shows the highest enrichment of pathogenic to benign variants to be near and within the most intolerant features defined by 3DTS. Shown are enrichment lines on a per Angstrom basis outside of the most intolerant site. The green line represents the enrichment of pathogenic missense variants over common (allele frequency > 1%) missense variants. Due to the scarcity of common variants, synonymous variants were also used to estimate neutral variation. The blue line indicates the enrichment of pathogenic missense variants over synonymous variants. Raw counts for each variant type with respect to distances are presented in Supplemental Figure S2. The reduction of number of counts at very close distances may be related to the spatial restrictions on distances smaller than a van der Waals contact.

Another application of the present work could involve prioritization of drug target sites. Protein structure-based methods are now routinely used at all stages of drug development, from target identification to lead optimization ^23^. Central to all structure-based discovery approaches is the knowledge of the 3D structure of the target protein or complex because the structure and dynamics of the target determine which ligands it binds ^23^. The characterization of human-specific intolerant sites and tolerance to genetic variation can be used to parse structural information to define active sites, but also to define functionally important topographically distinct sites that can support allosteric interactions. The presence of druggable, topographically distinct allosteric sites offers new paradigms for small molecules to modulate protein function ^24^. We analyzed the 3D intolerance characteristics for 102 proteins that included known drug targets with a bound ligand and proteins with known allosteric sites (**Supplementary Table S1 – available from authors**). The corresponding proteins carried a median number of one unique non-overlapping intolerant 3D-site (range 0-6). Overall, 18 proteins lacked an intolerant site, while 32 had greater than one unique intolerant site. Active sites were most constrained, followed by allosteric and ligand binding pockets (Figure 6A, Supplementary Figure S3). The lower scores of allosteric sites is consistent with the existing knowledge indicating that these sites tend to be under lower evolutionary conservation pressure than their orthosteric counterparts ^24^. We also observed an unequal distribution of tolerant and intolerant binding sites across therapeutic classes (Figure 6B). For example, antineoplastic and immunomodulating agents preferentially target intolerant sites. The identification of multiple intolerant 3D-sites and domains in many drug targets could be exploited for rational drug design and for analysis of drug screening results.

**Figure 6.**
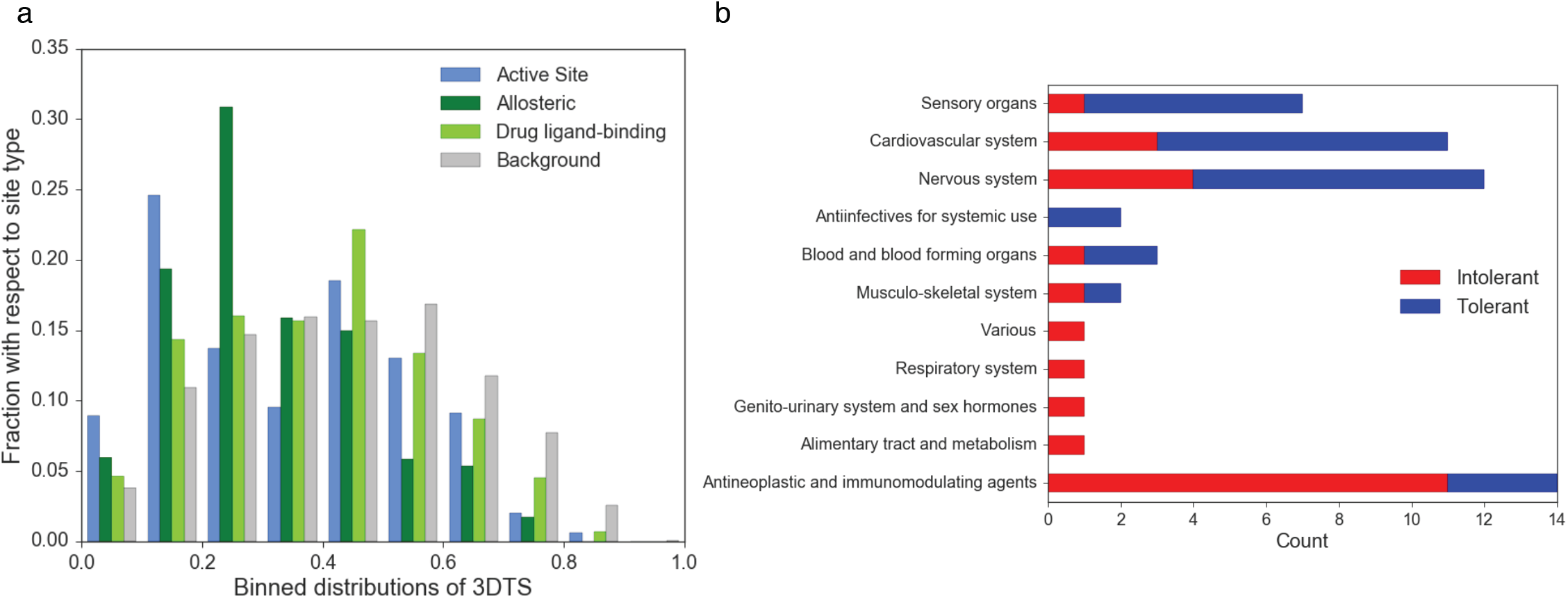
Characteristics of druggable sites. (a) Binned 3DTS scores describing active sites, allosteric sites, drug ligand-binding sites, and background. The sum of each site type is 1. Binned counts are provided in Supplementary Figure S3. (b) Counts of tolerant and intolerant drug ligand-binding sites grouped by therapeutic area. Here, tolerant is defined as 3DTS > 0.5, while intolerant is defined as described in the main text (3DTS < 0.33); drug binding sites between these 3DTS values are not included in (b). See Supplementary Table S1 for full details about this dataset.

In conclusion, the increasing detail of the limits of protein diversity that can be gathered through large scale sequencing of the human population provides valuable information on the functional consequences and pathogenicity of genetic variants and offers new data on orthosteric, allosteric and additional functional sites that can be harnessed for drug development.

## Methods

### Genomic and Variant Data

We included a set of 7,794 deep-sequenced unrelated whole human genomes (an extension from our previous report) ^2^, 123,136 exomes and 15,496 whole human genomes from gnomAD (http://gnomad.broadinstitute.org/). All data were aligned or lifted over to human reference genome assembly hg38. Variants from our set were included if they fell within the extended confidence region (as previously described) while gnomAD variant calls were included if they were annotated as “PASS”, and could be lifted over to hg38. Total call counts were derived from the sequence coverage files of gnomAD or from internal datasets. We used the transcripts and the gene models of Gencode version 26.

### Feature Data

Structural and functional feature annotations were taken from Uniprot text files (Downloaded 2017 April; www.uniprot.org ^25^) that were cross-referenced from Gencode. We used pairwise global sequence alignment to align the Uniprot amino acid sequence to the Gencode transcript sequence (after translating them to amino acids). The pairwise alignment algorithm was parameterized with the Blosum62 matrix, with a gap open penalty of 5 and a gap extension penalty of 1.

### Protein Structure Data Analyses

X-ray structure data from the Protein Data Bank (PDB; http://www.rcsb.org ^26^) were used if they were linked within the Uniprot text files. We used a pairwise global sequence alignment approach to align the Uniprot amino acid sequence to the amino sequence retrieved from the macromolecular Crystallographic Information File (mmCIF). Alignment parameters were set as above. The first author-defined biological assembly in the mmCIF file was used, when defined. In the cases in which it was not defined, the first biological assembly listed was used. In the case of the RING-domain of BRCA1, we used the NMR structure closest to the average as defined by the mmCIF file. The Pymol molecular visualization system (The PyMOL Molecular Graphics System, Version 1.8 Schrödinger, LLC.) was used to identify any residue within 5 Å of a defined Uniprot feature (also referred to as a “3D-site”).

### Quantification of Depletion of Variation and Creation of a 3D-Tolerance Score (3DTS)

Variation at genomic loci is modeled with independent Bernoulli trials. At loci i in individual j a variation happened with probability p. We assume that certain variants are incompatible with life in which case the variant is missing from the sample. Thus, the compound probability of observing a variant at a locus in an individual is *p*s*, where *p* is not specific to the variant, but is a genome wide mutation rate and s is specific to a variant with the interpretation of the probability that the variant is not lethal. If *s* = 0 then the genomic locus is completely depleted of variants, while if *s* = 1 then all variants are present as expected by the generic mutation rate of *p*. This simplistic model is not valid for common variants, however describes well the process of rare de novo mutation. In particular this model ignores: inheritance and relationship of individuals, linkage of variants by sharing a haplotype, allele frequency and zygosity. The main purpose of our work is to estimate the value of *s* because *s* is a good proxy of the strength of purifying selection on the genomic locus.

A nucleotide on the ancestral chromosome can change into three other nucleotides, not necessarily causing a non-synonymous mutation. We incorporate this into our model by extending it with the probability (b) that either of the three non-ancestral alleles lead to amino acid change. The value of *b* is derived from the genetic code and from the amino acid sequence of the transcript. With this extension, the probability of a mutation is *p * s * b*. To maximize power, we aggregate variants both by samples and by sets of loci induced by protein structure. Thus we write the probability of observing at least one variation at a given locus in *R* individuals: 1 – exp(- *p * s * b * R*). The latter follows from the Poisson approximation of the binomial distribution: the sum of the number of successes in *R* Bernoulli trials with the same parameter is a binomial distribution which can be well approximated with the Poisson distribution if *R* is large. The expression follows if we express ‘at least one’ as ‘not zero’.

To aggregate by different loci we treat each loci as a Bernoulli trial with parameter 1 – exp(- *p * s * b * R*). These parameters are however different for each loci, thus the sum of the number of successful trials is described with the Poisson binomial distribution. The expected value of the Poisson binomial distribution is the sum of its parameters (in our case sum(1 – exp(- *p * s * b * R*)), its density and distribution function may be approximated with the Poisson distribution (Le Cam’s theorem) or computed using Fourier transformations ^27^. For efficiency, we use the Poisson approximation.

Above we described the probability of observing at least one non-synonymous variant across a set of loci and a set of *R* samples assuming parameters *p* and *s*. We continue by estimating s from the available data assuming a global, genome wide constant value for the mutation rate parameter *p*. We estimate the constant mutation rate by numerically fitting the observed number of synonymous variants across all mapped proteins to the expected value of the number of synonymous variants fixing *s*=1. The estimated value is 2.5E-6 and from here on we treat it as constant. We calculate the posterior distribution of the remaining single s parameter by assuming a uniform prior between 0 and 1. In equations:

*P*(observed *k* variants in *K* loci among *R* samples | *s*) = Poi(k, sum over K(1 – exp(- *p* * *s * b * R*))), the likelihood function using Le Cam’s approximation

*P*(*s*) = 1, uniform prior over 0-1

*P*(*s* | observed *k* variants in *K* locus among *R* samples) = Lik * prior / integrate over 0-1 (lik*prior), the posterior

We summarize the posterior distribution of *s* by its mean value, which we assign to each protein feature and refer to it as 3D tolerance score (3DTS) elsewhere in the paper.

### Functional Data and Pathogenicity Scores

Functional *in vitro* data for PPARG were sourced from Majithia et. al.^19^ The integrated functional scores available through http://miter.broadinstitute.org/ (Data version 1.0) were used. Only those scores linked to amino acid changes resulting from a single nucleotide variation were used in the comparison with 3DTS.

Functional *in vitro* data for the RING domain of BRCA1 were sourced from Starita et. al.^20^ Known homology directed repair (HDR) rescue scores from the HDR rescue assay were used when available, otherwise predicted values were used. Only those scores linked to amino acid changes resulting from a single nucleotide variation were used in the comparison with 3DTS.

### Variant Distance Data and Analyses

Distance-based quantification was performed using Pymol. Pathogenic variation data was sourced from Clinvar ^28^ (July 2016) and HGMD ^29^ (first quarter 2016, R1). Selected Clinvar variants had to be tagged as (likely-)pathogenic and have 1 or more stars. Selected HGMD variants had to be tagged as DM and High. Any pathogenic variants overlapping a variant annotated as benign with 1 or more stars in Clinvar were filtered out. Structures were included in this analysis if at least 80% of the total canonical protein length was covered and at least one pathogenic missense variant was present.

### Drug Ligand Data Set and Analyses

A set of structures defined as therapeutic targets of FDA-approved drugs was used. Therapeutic targets were taken from the supplementary information of Santos et al.^30^ Of 667 non-redundant Uniprot entries, 361 contained some structural information and 100 contained proteins where the sequence length of the structure defined by Uniprot covered at least 80% of the canonical Uniprot sequence. Ninety-four of these 100 proteins were mapped to the genome using Gencode version 26. These 94 proteins were examined for the presence of the corresponding bound therapeutic molecule or analog in a structure; when not found, homologous structures containing these molecules were superimposed, resulting in 48 structures with their corresponding “bound” therapeutic molecule (for a list of these structures and their “bound” ligands, see **Supplementary Table S1**). Ligand binding sites were defined as those residues within 5 Å of any of the bound therapeutic molecule residues. The lowest 3DTS value was assigned to each of these residues in cases of overlapping 3D-sites.

### Anatomical Therapeutic Chemical (ATC) Classification System Data and Analyses

Drug-liganded molecules (as identified in the above Drug Ligand Data Analyses section) were assigned to their ATC codes using the supplementary information of Santos et al.^30^ For each structure, a non-redundant list of top-level ATC code was included for all bound drugs. In cases in which no ATC code was found, the code was inferred either based on indication (when available) or based on indirect effect. In cases where the structure had multiple chains contributing to the ligand-binding site, the median score was used in defining tolerance.

### Allosteric Data Set and Analyses

The XML data of the Allosteric Database (Release 3.06)^31^ was downloaded and parsed with custom Python scripts. Data was used if the field “Organism_Latin” was equal to “Homo sapiens”, any of the allosteric counts (“Allosteric_Activator_Count”, “Allosteric_Inhibitor_Count”, or “Allosteric_Regulator_Count”) had a value of at least one, and “Site_Detail” contained at least one defined amino acid. Of the resultant fifty-four entries, fifty structures were mapped where every allosteric residue had a 3DTS value. The lowest 3DTS value was assigned in cases of overlapping 3D-sites. These structures were used in the downstream analysis (for a list of these structures, see Supplementary Table S1).

### Active Site Data Set and Analyses

A non-redundant list of protein active sites was included for those structures found in the Drug Ligand Data Set and the Allosteric Data Set. Active sites were defined based on the 5 Å context of the “ACT_SITE” feature(s) defined in Uniprot (i.e., “ACT_SITE” 3D-sites).

### Unique, Non-Overlapping 3D-Intolerant Site Analyses

Structures from the Drug Ligand Data Set and Allosteric Data Set were used. A 3D-site was defined as intolerant if the 3DTS value was in the 20^th^ percentile proteome-wide (3DTS value < 0.33). 3D-intolerant sites were joined if at least one residue overlapped within a chain. For homomeric chains, two intolerant sites were considered unique if no residue in the primary structure was shared. In cases where chains representing the same protein differed in the number of unique, non-overlapping 3D-intolerant sites, the maximum number of 3D- intolerant sites was chosen.

### Statistics

Plots were produced using the Seaborn (https://seaborn.pydata.org) and Matplotlib (https://matplotlib.org) libraries in Python. Statistics were calculated using the Numpy (http://www.numpy.org) and SciPy (http://www.scipy.org/www.scipy.org) libraries in Python and in house statistical software in Scala.

## Acknowledgments

The authors would like to thank the Genome Aggregation Database (gnomAD) and the groups that provided exome and genome variant data to this resource. A full list of contributing groups can be found at http://gnomad.broadinstitute.org/about.

**Author contributions**. Conception and design of the study: M.H, I.B., J.C.V., A.T. Performed the analyses: M.H., I.B., J.diI., R.A. Wrote the manuscript: all authors.

## Supplementary Materials

**Supplementary Table S1.** Data used in the construction of Figure 6.

**Supplementary Figure S1**. Correlation between in vitro functional data and 3D-site in tolerance for PPARG.

**Supplementary Figure S2**. Relationship between tolerance and pathogenicity.

**Supplementary Figure S3**. Characteristics of druggable sites.

## Supplementary Figures

**Supplementary Figure S1.**
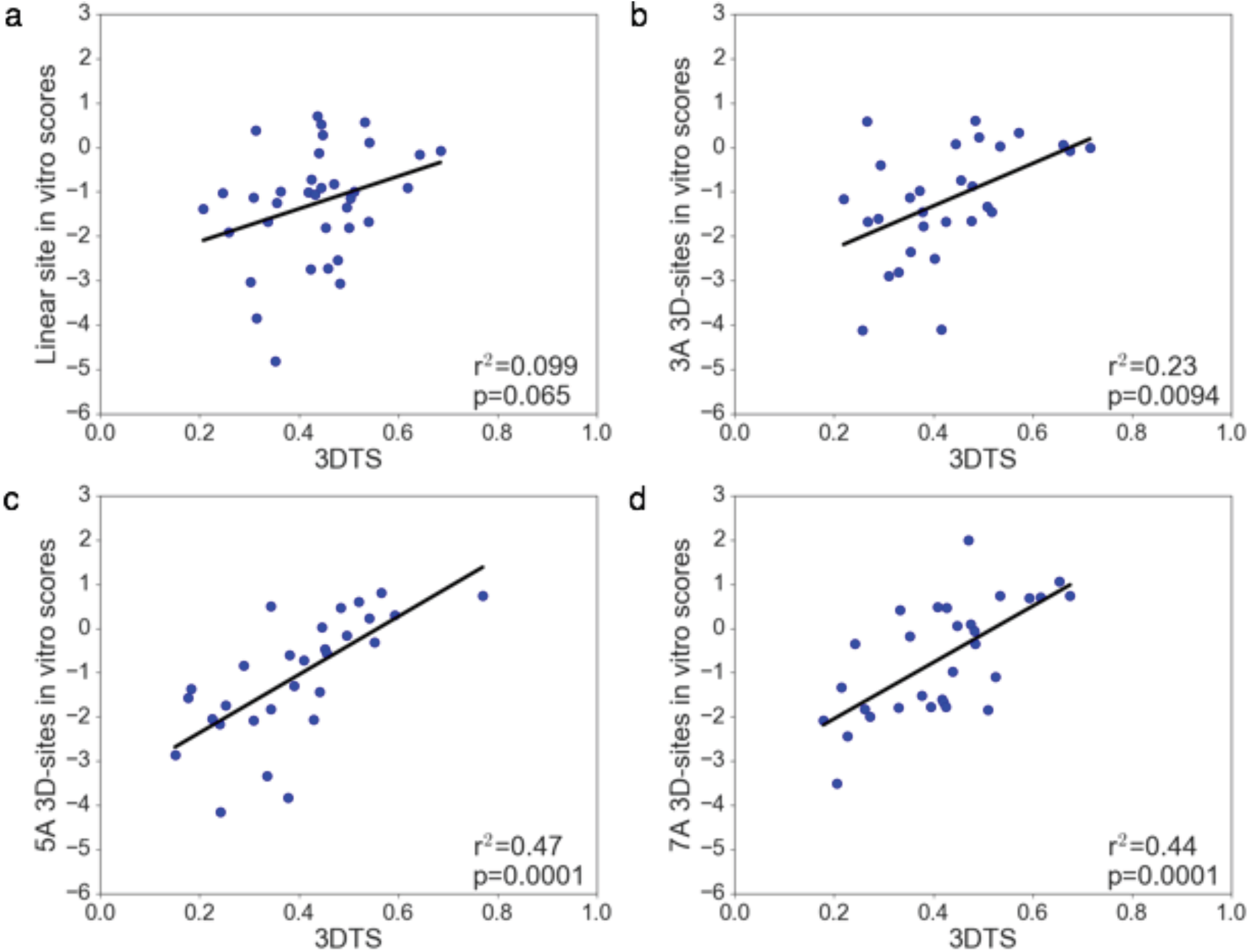
**Correlation between in vitro functional data and 3D-site in tolerance for PPARG.** The 5 Å 3D site (r^2^=0.47; Figure 3) performs best relative to a linear site-approach as well as other 3D distances in a correlation analysis with *in vitro* data for PPARG. The distances tested included (a) the linear site (no 3D context added; r^2^=0.099), (b) 3 Å (r^2^=0.23), (c) 5 Å (r^2^=0.47), and (d) 7 Å (r^2^=0.44).

**Supplementary Figure S2.**
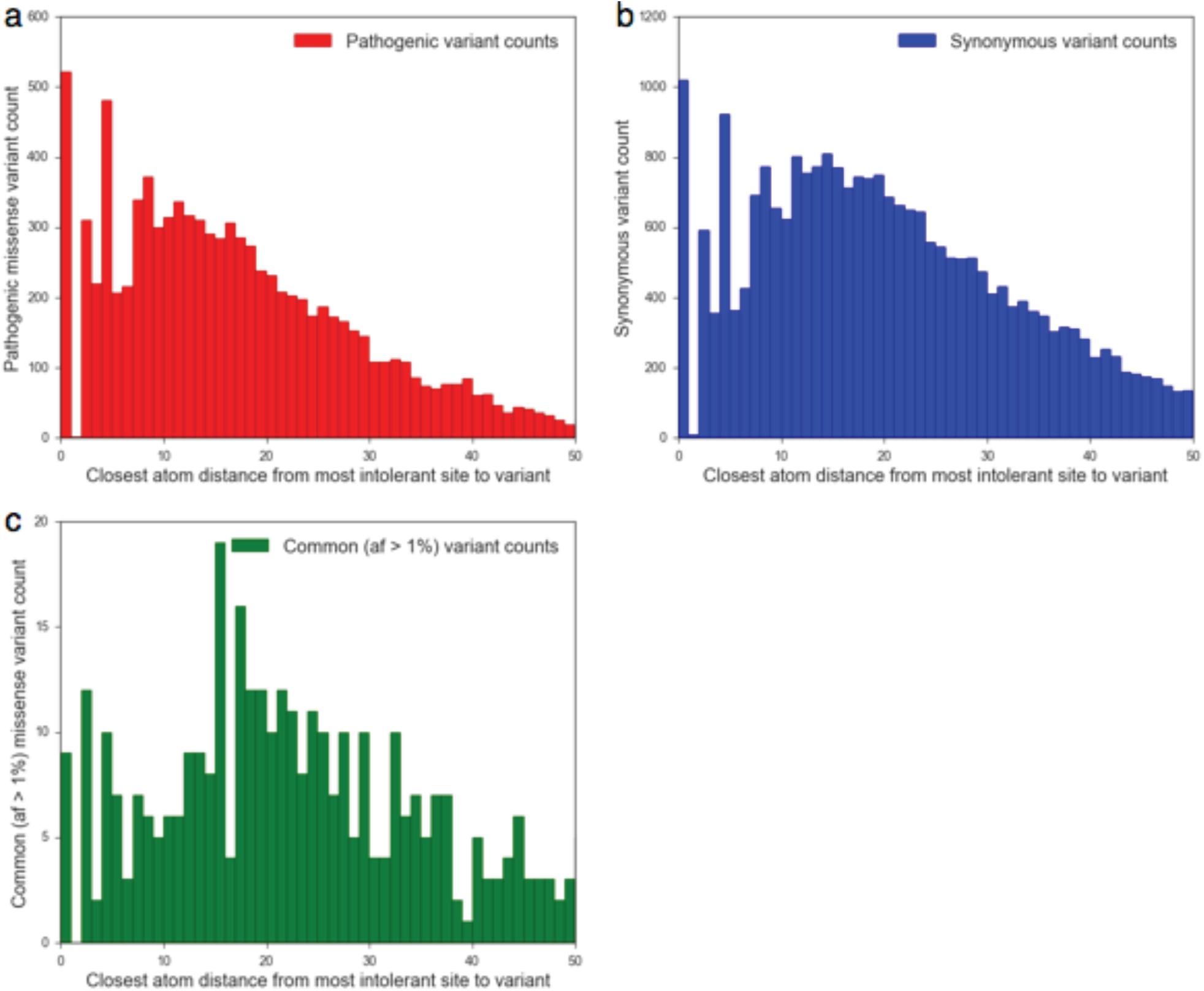
**Relationship between tolerance and pathogenicity.** Raw counts and distances corresponding to Figure 5 for (a) pathogenic missense variants, (b) synonymous variants, and (c) common (allele frequency > 1%) missense variants. Note that the first bin represents the information within the most intolerant 3D site and subsequent bins represent the counts only within each binned distance. The apparent “noisy” first few bins are due to biophysical constraints placed on inter-residue interactions (i.e., a minimum of about two Angstroms to find additional residues followed by more distance to identify subsequent residues, dependent upon orientation and residue types). The reader is directed to Tables II and III in Seeliger and de Groot, 2007 ^33^ for additional background on atomic radii and lower bounds for distances of specific atom type combinations in proteins.

**Supplementary Figure S3.**
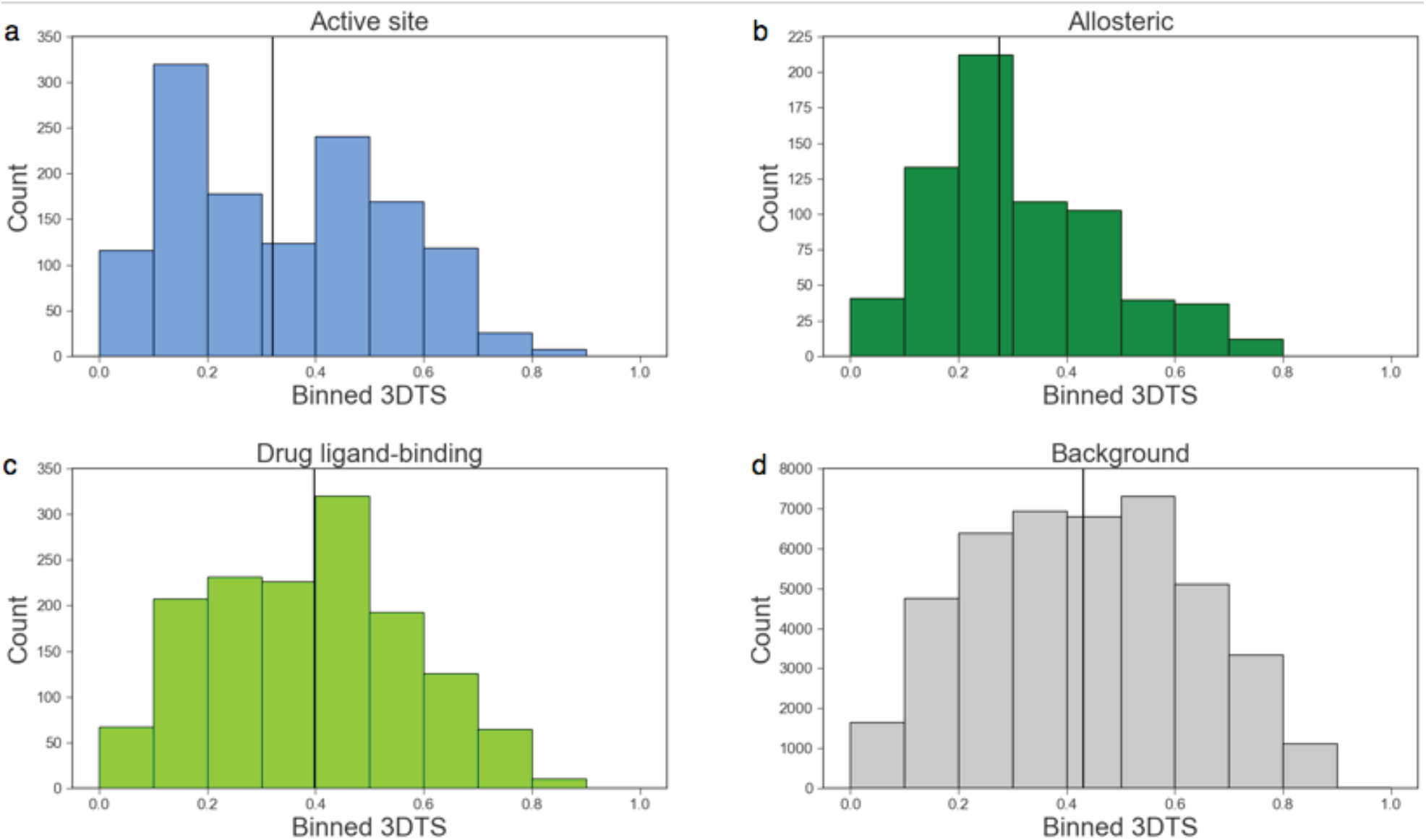
**Characteristics of druggable sites**. Histograms corresponding to the data presented in Figure 6A for (a) active sites, (b) allosteric sites, (c) drug ligand-binding sites, and (d) background. Colors are as in Figure 6A. The median of each plot is drawn as a vertical line.

